# Correcting machine learning models using calibrated ensembles with ‘mlensemble’

**DOI:** 10.1101/2021.07.26.453832

**Authors:** Tomasz Konopka

## Abstract

Machine learning models in bioinformatics are often trained and used within the scope of a single project, but some models are also reused across projects and deployed in translational settings. Over time, trained models may turn out to be maladjusted to the properties of new data. This creates the need to improve their performance under various constraints. This work explores correcting models without retraining from scratch and without accessing the original training data. It uses a taxonomy of strategies to guide the development of a software package, ‘mlensemble’. Key features include joining heterogeneous models into ensembles and calibrating ensembles to the properties of new data. These are well-established techniques but are often hidden within more complex tools. By exposing them to the application level, the package enables analysts to use expert knowledge to adjust models whenever needed. Calculations with imaging data show benefits when the noise characteristics of the training and the application datasets differ. An example using genomic single-cell data demonstrates model portability despite batch effects. The generality of the framework makes it applicable also in other subject domains.

## Introduction

Machine learning (ML) models are routinely used for analyzing biological data for research purposes (Camacho et al. 2018) and are starting to be deployed in translational settings (Wiens et al. 2019; Kann et al. 2021). The adoption of ML is in part forced by the increasing volumes of data that cannot be processed manually, and it can also be justified by promising performance. However, intangible factors also play important roles in how ML can influence a research domain (Gunning et al. 2019). The need to understand how automated systems make predictions, for example, has spurred progress on conceptual (Lipton 2018) and practical fronts (Samek et al. 2019). As a result, several tools have been developed to explain how ML models reach predictions (Molnar 2019; Biecek and Burzykowski 2021). Other aspects that can build confidence are rigorous testing in appropriate translational settings (Wiens et al. 2019) and the possibility to update imperfect models (Babic et al. 2019; Wu et al. 2021). While testing is an ongoing and subject-specific process, the task of updating can be approached in a systematic fashion. This work explores strategies to adjust existing ML models to improve their suitability and performance on new data. It also offers a practical software implementation.

In a canonical machine learning study, a model is trained on a tranche of an initial dataset and evaluated on a separate tranche. The model can then be deployed under the assumption that new data will be similar to the training set. Over time, however, incentives may appear to change the model. For example, a sudden shift or a gradual drift in the properties of incoming data may blunt model performance. Richer incoming data may raise expectations that the original model cannot meet. Identification of a bias in the initial dataset may prompt the requirement to correct improper predictions (Jiang and Nachum 2019).

Incentives to update a model can also arise in biomedical settings. A model may require adjustment if it is trained at one laboratory but later deployed at another site where, despite best efforts for consistent protocols, measurements are performed with different batches of reagents or environmental conditions (Wu et al 2021). The number of data features can change if a laboratory seeks to improve predictions through an integrative approach with auxiliary measurements. Finally, it is possible that biases may be detected in bioinformatic models. Databases such as the UK Biobank (Sudlow et al. 2015), for example, are known to hold a skewed representation of populations (Fry et al. 2017). While some findings are generalizable to wider populations (Batty et al. 2019), the need to update ML models may arise in the future, for example, in the context of sub-populations or in light of changing environmental factors.

A direct solution to such situations is to train a new model on a combination of the original and new data. After testing the new model on independent datasets, it can be deployed as a replacement of the original model. In principle, this strategy should yield a model that incorporates existing strengths and meets any new requirements. However, this direct approach may not be feasible in practice. For example, the original training data may no longer be accessible because of privacy restrictions. Thus, this work explores possibilities for correcting existing models without re-training from scratch. It considers tasks where the inputs consist of a pre-trained model and a new labeled dataset, but not the original training dataset.

This setting bears similarity to transfer learning in that the intention is to use existing knowledge to create a new model (Zhuang et al. 2021). However, transfer learning methods require substantial retraining and in some cases rely on in-depth knowledge of the internal model architecture, for example interconnections in neural networks. Such methods are not necessarily straightforward to use on a routine basis. In other contexts, researchers enhance predictions by polling multiple alternative methods and producing a consensus. This approach has been used in cancer genomics for pattern discovery (Guinney et al. 2015; Dentro et al. 2021) and classification (Ellrott et al. 2018). However, projects have relied on manual curation of the constituent algorithms and pooling protocols. Tools are thus needed to make such methodology more accessible in common practice.

The main contributions in this work are threefold. First, a taxonomy for correcting machine learning models is introduced. This shadows categories already employed for grouping algorithms in terms of pre-processing, post-processing, and calibration (Jiang and Nachum 2019), and incorporates the concept of model ensembles (Caruana et al. 2004). The taxonomy serves to elucidate the problem at hand and helps to approach it in a systematic fashion.

Second, the taxonomy guides the design of a software package ‘mlensemble’ for building and calibrating ensembles of machine learning models in the R environment (R Core Team 2021). The package overlaps in some features with ‘caretEnsemble’ (Deane-Mayer and Knowles 2019). However, while ‘caretEnsemble’ creates ensembles by training on a common dataset, ‘mlensemble’ is designed for heterogeneous models, including models trained on different datasets. The techniques implemented in the package have all been employed in bioinformatic studies either as part of advanced ML algorithms or in project-specific methods. The contribution of the package is to make ensemble creation and calibration accessible through a few commands in the R environment.

Third, a series of examples using image and genomic datasets show the possibilities for correcting ML models in practice. Explorations with the MNIST digits dataset illustrate how the performance of reasonable models can drop when the characteristics of new data deviate from those seen during training, and how such effects can be attenuated with the ‘mlensemble’ package. Explorations with single-cell transcriptomic data translate the concepts to genomics. They show how trained models can be reused across datasets, even in the presence of biological and technical shifts.

## Results

### Strategies for correcting machine learning models

A typical study in supervised machine learning starts with a raw dataset and associated labels (continuous or discrete) and trains a model, which can then make predictions on new data (Figure 1a). After a model is finalized, there may appear incentives to update it. While details are bound to be application-specific, it is possible to categorize the possibilities in an attempt to formulate a taxonomy of strategies for correcting ML models.

**Figure 1.**
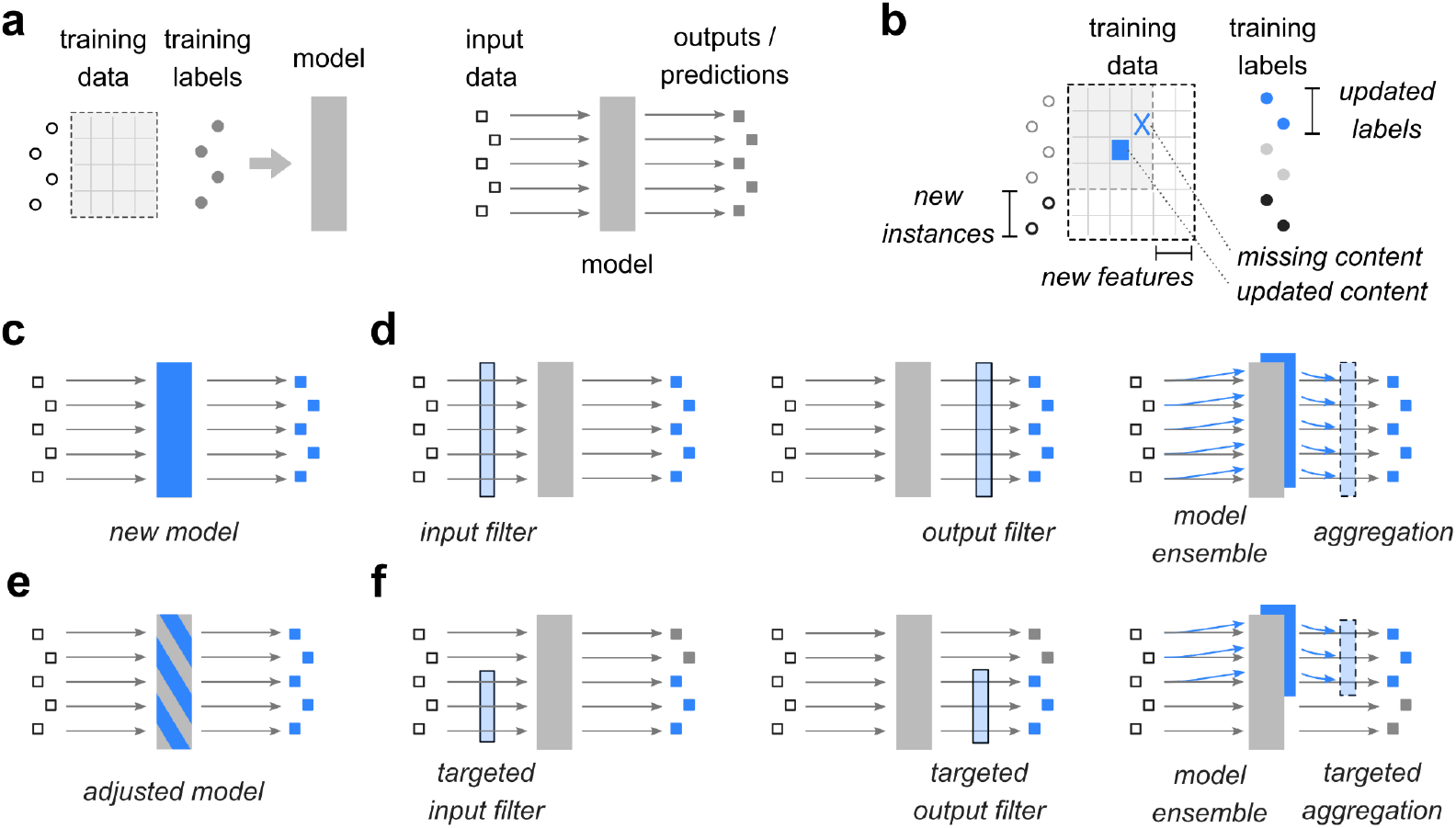
Strategies for correcting machine learning models. **(a)** Schematic of a typical machine-learning study with a training dataset and training labels, which can be used to construct a model. The model can be used in production on new input data. **(b)** Schematic of changes that may require adjusting or correcting an already-trained model. **(c)** Replacing the original model by a new model constitutes a global strategy for adjusting an existing workflow. **(d)** Other global strategies for adjusting an existing workflow through pre-processing, post-processing, and use of multiple models in parallel. **(e**,**f)** Analogous to (c,d), but showing targeted strategies that affect small portions of the workflow components.

Incentives to update a model stem from changes in one of the components used to build the model (Figure 1b). Availability of new training data items can raise the expectation that an updated model may perform better than the original. New features can create opportunities to find more associations between raw data and output labels. Inaccurate content in the dataset can be corrected, leading to data entries changing values, missing entries being filled in, and output labels changing. Overall, these scenarios can be categorized as changes in the data instances, features, content, and expected outcomes.

Machine-learning studies can respond to a changing environment in many ways. One way to categorize possible approaches is to label them as global (Figure 1c,d) or targeted (Figure 1e,f). The defining characteristic of a global adjustment is that it affects the machine-learning logic in a uniform manner. The most radical global adjustment is to replace an original model by a new one (Figure 1c). Other global strategies that preserve a role for the original model include pre-processing of input data, post-processing of model outputs, or establishing an ensemble of models (Figure 1d). An example of a relevant pre-processing step is imputation. This can take the form of applying simple recipes to replace missing data by placeholder values, but it can also be a directed process that corrects specific biases (Konopka et al. 2021).

Targeted strategies can be defined in analogy to the global ones, but where adjustments are made to small parts. The analog to replacing the entire model is tuning existing components within the original model (Figure 1e). In an intrinsically interpretable model such as a linear regression, this can, for example, entail changing fitted coefficients. This strategy encompsses transfer learning (Zhuang et al. 2021) and lifelong learning (Zhang et al. 2019). However, these approaches require detailed insight into how a particular model functions. They are therefore not easily generalizable and are not covered in this work. By contrast, analogs to global pre-processing, post-processing, or ensemble approaches can be applied to arbitrary models, even those that are not intrinsically transparent (Figure 1f).

The distinction between global and targeted strategies is blurred. In particular, a targeted pre-processing step can always be implemented as a uniform pre-processing filter that internally adjusts some inputs and leaves others unaffected. The same applies to post-processing and ensemble strategies. Nonetheless, the taxonomy provides a language to convey the possible ways by which a machine-learning model can be augmented or adjusted.

A framework that mirrors the taxonomy is implemented in an R package, ‘mlensemble’ (Methods, Supplementary Table 1). Importantly, the package can manage multiple models trained on disparate datasets, and aggregate their predictions into an integrated result. The package can also calibrate models and ensembles. This is a form of post-processing that uses additional data items to determine how the aggregation should be weighted to reproduce expected outcomes.

### Correcting multiclass classification models

To demonstrate correcting ML models in practice, it is instructive to apply the framework on a dataset that is neutral with regard to subject domain. The MNIST Digits dataset, which consists of 70,000 images of handwritten digits (LeCun et al. 1998), is convenient as it is also readily visualized. To begin experiments with this data, images were split into disjoint sets for training (40,000 items), holdout (10,000 items), and testing (remaining 20,000 items). Next, a series of dataset pairs were created (Figure 2a) with the intention that the first dataset in each pair would be used to train a primary model and the second dataset and any holdout tranches would be for correcting.

**Figure 2.**
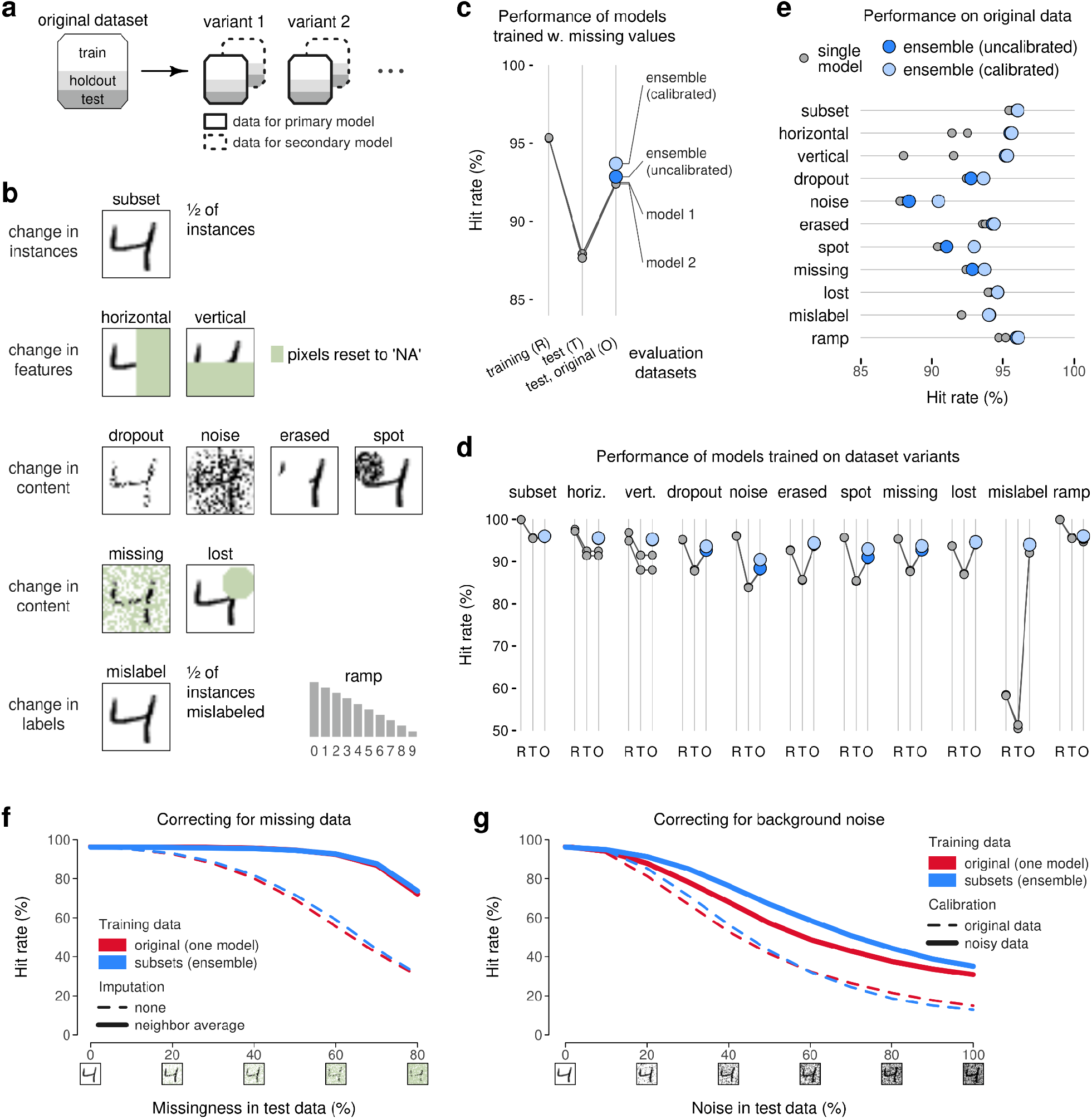
Correcting multiclass classifiers. **(a)** Schematic of partitioning a dataset into sections for training, holdout, and testing. The original dataset is used to create variants, which are split into primary and secondary datasets. **(b)** Schematic of variants of the MNIST digits dataset; images show how a representative image is mutated in each variant. **(c)** Classification performance obtained by models trained on an MNIST variant with missing pixels. Performance is evaluated on the training images (R), test images with missing pixels (T), and test images from the original dataset (O). **(d)** Extension of (c) to all dataset variants. **(e)** Alternative summary of models trained on MNIST variants and evaluated on test images of the original dataset. **(f)** Classification performance of a model and an ensemble on test images with increasing amounts of missing pixels. Performance is evaluated with and without a pre-processing filter that imputes missing values by averaging neighboring pixels. **(g)** Classification performance of a model and an ensemble on test images with increasing amounts of background noise. Performance is evaluated using models calibrated on noise-free data, and with separate calibration at each noise level.

Dataset were created to simulate a range of scenarios (Figure 2b, Methods). For example, one variant denoted as ‘subset’ separated the original images into primary and secondary datasets at random, producing datasets that were structurally similar to the original, except smaller in size. They represent a scenario where data are collected at multiple sites with identical protocols. Other variants simulated changes in the available features, changes in content, and changes in the labels (Figure 2b, Methods).

Separate multiclass classification models were trained on the original dataset and on the primary and secondary datasets for each variant. For simplicity and transparency, all the models were trained in the same way with the same hyperparameters (Methods). The hit-rate (equivalent to recall, or the fraction of images in the test set labeled with the correct digit) for the model trained on the original dataset was 96.1%. For the current purpose, it is not important that this is not the maximal achievable performance. The result merely defines a level of performance that is achievable given the resources available for training and given access to the entire (training) dataset.

As expected, models trained on two portions of the variants performed less well than the baseline model. For example, models trained on primary and secondary datasets of the variant with missing values achieved a 95.3-95.4% hit rate on their training images and 87.7-87.9% hit rate on their test images (Figure 2c). The objective of the exploration, however, was to improve performance of the primary model given the possibility to use the primary model together with additional information. Two ensembles were thus constructed. One ensemble simply averaged predictions from models trained on the primary and secondary datasets. In the second, the contributions of the two individual models were calibrated using the holdout tranche of the original dataset. The uncalibrated and calibrated models achieved hit rates of 92.9% and 93.7%, respectively, thus outperforming either one of the individual models (Figure 2c).

Similar patterns were observed for all the dataset variants (Figure 2d). Interestingly, the performance of models trained on the ‘mislabel’ variant with a high fraction of erroneous labels was very poor on test images which were themselves mislabeled. However, performance of the ensembles rose to a comparable level to other ensembles when evaluated on test images with correct labels. This is an example of how performance can appear to rise in a test environment compared to a training environment if the quality of data in the former is higher than in the latter.

Focusing on the performance of individual models and ensembles on the original test images, the overall results indicated a consistent benefit to using ensembles (Figure 2e). For some variants, the ensembles approached the baseline performance of the model trained on the original data.

While ensemble-building is a strategy that can be applied in almost all situations, some variants could also be approached with alternative and bespoke pipelines. For example, imputing pixel intensities is a reasonable strategy for datasets with missing values. To explore this, a series of datasets were created with increasing levels of missingness (Methods). As expected, model performance decreased with the level of missingness (Figure 2f), both for an individual model and for an ensemble. Imputing missing values through a convolution extended the utility of both those approaches. This demonstrates the scale of the discrepancies that may arise from using the same models in pipelines with and without appropriate preprocessing.

A related result was observed in simulations where images were subjected to increasing levels of noise (Methods, Figure 2g). Performance of an individual model and a model ensemble decayed with the level of noise, as expected. However, there was a noticeable effect due to calibration. When the models were calibrated using holdout images from the original dataset (i.e. without noise), the individual model and the model ensemble performed comparably. When calibration was performed using holdout images with noise, the overall performance increased at all noise levels. In addition, the ensemble consistently outperformed the single model. More in-depth inspection of the predictions revealed that noise levels triggered an imbalance in the predicted labels, which could be alleviated to some extent via calibrations (Supplementary Figure S2). These results not only highlight that an existing model can be corrected or improved, but also highlight the need for calibration before applying a model to new data with different noise characteristics.

### Correcting models in genomics

In genomics, measurements of RNA from single cells produce gene-expression profiles for thousands of cells, making machine-learning methods essential for analysis (Kharchenko 2021). Such measurements provide insights, for example, about the diversity of cell types within a tissue and about organ development. Gene-expression data are also known to exhibit batch effects, making it challenging to generalize findings from one dataset to another. Indeed, reliable classification of cell types using expression of marker genes is typically achievable only upon integrating data from several training datasets (Fischer and Gillis 2021). This leads to the question of how to correct a cell-type classification scheme devised from an initial exploratory study to help analyze new studies.

To explore correcting gene-expression models, two datasets were selected based on their shared goal to characterize cell types (Methods). The first dataset consisted of over 50,000 profiles from cells from the testes of two adult donors (Sohni et al. 2019). The original study characterized the gene expression profiles of several cell types (Sertoli cells, Leydig cells, peritubular myoid, endothelial cells, and others), and published sets of marker genes. The 103 marker genes can be used to visualize the dataset in low dimensions and gene expression for cell-type specific marker genes can act as a simple cell-type classification scheme (Figure 3a, Methods).

**Figure 3.**
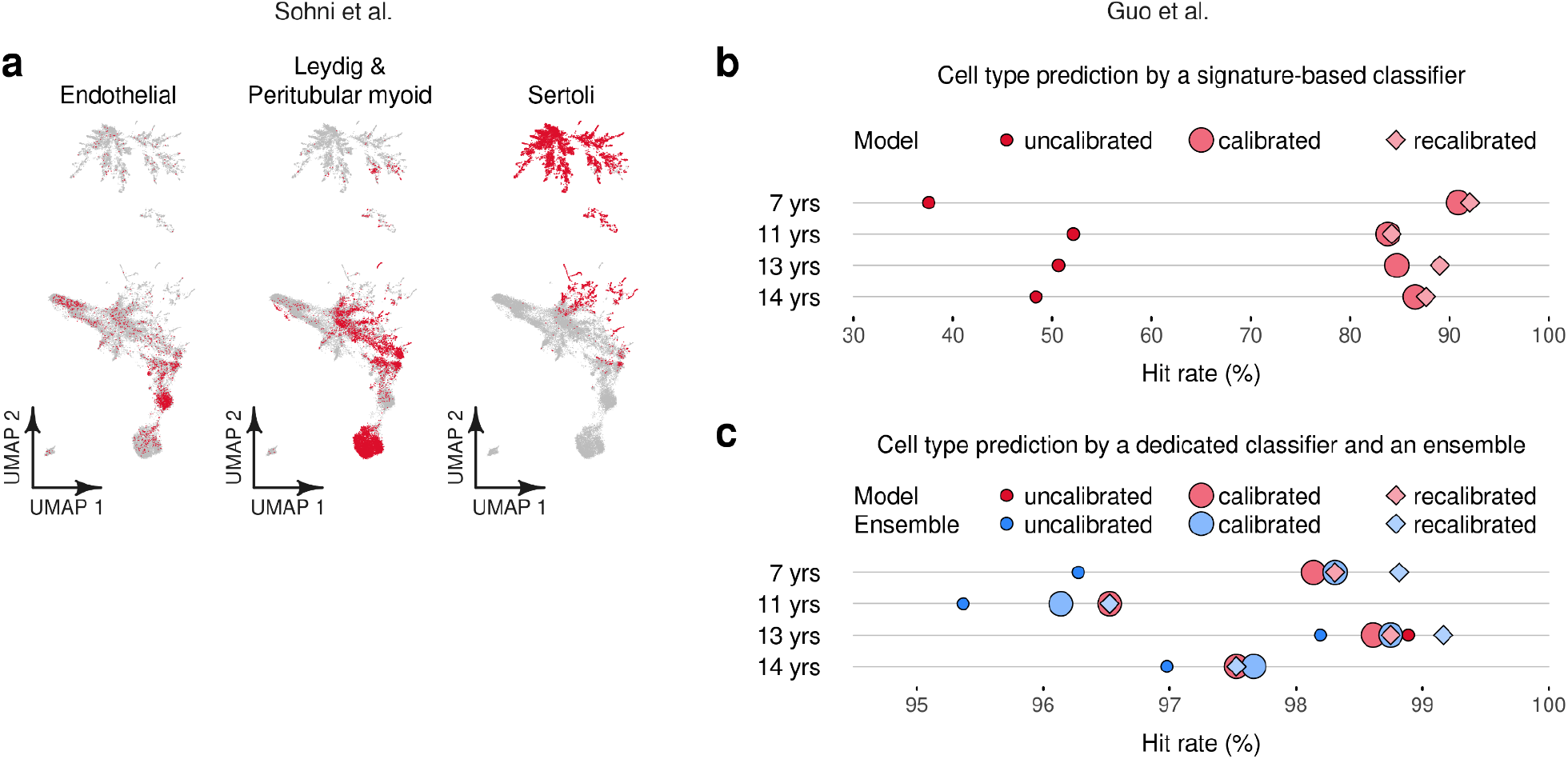
Correcting cell-type classification. **(a)** Embeddings of gene-expression profiles from single cells from testis. Dots represent individual cells. Embeddings were generated using gene-expression values for 103 marker genes. Panels highlight selected cell types in color; cell types are determined by the dominant gene expression signature. **(b)** Performance of cell-type prediction in an independent dataset. Performance was measured for a signature-based classifier without calibration, with calibration to the entire new dataset, and with recalibration to sample-specific data. **(c)** Analogous to (b), but with predictions performed using an xgboost multiclass classifier and an ensemble of the xgboost and the signature-based classifiers.

The second dataset consisted of over 31,000 single-cell profiles from the testes of six donors, of which 7,413 cells from four donors were labeled with a cell type (Guo et al. 2020). The labeled cells in this dataset offer an opportunity to explore how gene signature classifiers defined in one study can reproduce independent labels in a second dataset. To this end, the second dataset was restricted to the same 103 marker genes and split into portions for training, holdout, and testing. Because of batch effects and differences in donor age across the two datasets, direct application of a classifier on new data was expected to yield unremarkable performance. Indeed, the rate of correct cell-type classification by the gene signature-based classifier on the test tranche of the second dataset hovered around 50%, with variability when stratified by donor (Figure 3b). However, calibration of the signature-based classifier using holdout data increased classification performance to over 80%. Recalibration to donor-specific holdout data improved performance further. These results demonstrate that using a fraction of data from a new dataset for calibration can yield a substantial enhancement to the ability to predict cell types, even in the presence of technical and biological effects.

Given labels for all cells, it is possible to train a dedicated classifier on the new data, as well as to use the dedicated classifier together with the initial gene signature-based approach (Methods). A dedicated classifier trained on a specific dataset has the advantage that it is by construction adjusted to the peculiarities of the data. Experiments involving a dedicated classifier achieved cell-type prediction performance in excess of 95% (Figure 3c). Interestingly, the performance of the dedicated classifier changed little when it was trained and applied without modification to the test data, calibrated with the entire holdout set, or recalibrated with donor-specific data (Figure 3c).

Using a dedicated classifier in an ensemble with the initial gene signature-based classifier produced more variability. The uncalibrated ensemble received the lowest performance scores - lower than the dedicated classifier alone. This was expected as incorporating a poor predictor in a naive ensemble is bound to produce poor outcomes. However, calibration restored competitiveness and recalibration on donor-specific holdout sets resulted in the best overall performance in three out of four samples (Figure 3c). This demonstrates that incorporating prior knowledge in the form of a pre-trained classifier can enhance performance even of a dedicated classifier, as long as the ensemble is calibrated before performing predictions on new data.

## Discussion

Machine learning techniques are used routinely in biology research, for example in genomics, but most projects train models from scratch, and model reuse is rare. With the increasing availability of reference datasets and with increasing sizes of primary datasets, creating models from scratch is set to become proportionally more time-consuming. This points to transfer learning as a technique to streamline the deployment of ML for new applications. This work addressed the scenario wherein a trained model exists for a task, but should be updated to correct imperfections or tune performance to a new setting. It defined a taxonomy for correcting ML models, presented a software implementation, and presented a series of examples using general-interest and genomic datasets.

The taxonomy defines building blocks for correcting ML models. These building blocks shadow steps already used in bespoke ML pipelines as well as frameworks for automatic model creation (Zoller and Huber 2021). Their purpose here is to guide development of a practical software package, ‘mlensemble’. The systematic nature of the taxonomy conveys that the package offers a diverse set of options for correcting ML models. At the same time, the taxonomy is not necessarily complete; it would be natural to extend it, for example, through nested structures. Any extensions can guide the integration of additional features into the package.

Experiments with variants of MNIST digits and with genomic single-cell expression datasets demonstrate that the ‘mlensemble’ package is applicable in a range of applications. Its ability to construct and calibrate ensembles can bring substantial benefits over naive methods. In particular, calibration reduced class imbalance in the presence of noise (Figure 2g, Supplementary Figure 2). It also enhanced the performance of a cell-type classifier based on gene signatures from around 50% to over 80% when deployed on a new dataset (Figure 3b). Such benefits of ensembles and calibration are not surprising (Caruana et al. 2004). However, while these techniques have been utilized internally by various ML algorithms, ‘mlensemble’ exposes these techniques to the application level. It enables analysts to join models into ensembles in a manual manner, yet with a straightforward interface. This approach is consistent with how ensembles are used in bioinformatics, i.e. joining complete and well-motivated models to increase robustness (Guinney et al. 2015; Ellrott et al. 2018; Dentro et al. 2021).

Construction of simple ensembles with ‘mlensemble’ opens opportunities to reuse working models and fine-tune them with a fraction of the data required to train a new model from scratch. In genomics, for example, this gives analysts the option to label only a fraction of items for calibration to achieve moderated performance (Figure 3b) instead of labeling a very large number of data items to train and validate a new dedicated model (Figure 3c). Even in the event of the latter, this approach offers a fast method to integrate prior knowledge.

It should be noted that the approach for correcting ML models implemented in ‘mlensemble’ does not give formal guarantees. Performance gains are moreover dependent on the quality of the data used for calibration. Thus, the package does not remove the need for rigorous assessment of the updated ML models, which should be carried out using established methods.

In summary, the software package provides tools for correcting ML models that can be employed in everyday analyses. The package should facilitate model sharing and reuse.

## Methods

### R package ‘mlensemble’

A framework for ensemble building was implemented as a package for the R environment (R Core Team 2021). The package introduces two main types of data objects, accompanying functions, and some helper objects.

The first data object type is ’ml_model’. Objects of this type wrap R models or functions. Because of this design, the framework can support a wide range of models created using base R, third-party packages like xgboost (Chen et al. 2019), and, in principle, also models implemented outside of the R environment. The framework supports regression as well as multi-class classification. ‘ml_model’ objects also support custom hook functions for pre-processing and post-processing data.

The second data object type is ‘ml_ensemble’, which contains a list of ‘ml_model’ objects. The purpose of this object type is to manage ensemble constituents. Objects can be constructed by joining several ‘ml_model’ objects together using the ‘+’ operator.

The package implements the `predict` method for ‘ml_model’ and ‘ml_ensemble’ objects. Thus, these objects can be used within workflows in the same way as other models. Predictions automatically apply pre-processing and post-processing steps. In the case of ensembles, predictions also harmonize the outputs from constituent models. Thus, if models create different label sets or provide labels in different order, such discrepancies can be resolved automatically.

Importantly, a function `calibrate` allows users to tune an ensemble to a specific use-case or type of dataset. For regression tasks, calibration trains a single linear regression model using predictions from constituent models. During prediction, this model uses learned weights to produce a joint prediction. For classification tasks, a separate logistic regression model is trained for each output class. During prediction, these models make adjustments to the probabilities associated with each class. Because the transformations are independent, they do not guarantee that the adjusted probabilities add up to unity across all the classes. Thus, multi-class ensembles also carry out an additional normalization step.

The calibration procedure is a straightforward application of linear and logistic regressions. They are therefore fast to train and to use. The assumption that multi-class calibration can be performed on each label independently is a strong simplification. However, it can have practical utility when the aim is to introduce mild adjustments to well-performing models. The value of the ‘mlensemble’ package lies in a coherent interface to manage pre- and post-processing, ensemble methods, and calibration methods (Supplementary Table 1).

### MNIST digits dataset and variants

The MNIST digits dataset was downloaded from http://yann.lecun.com/exdb/mnist/ and partitioned into a training set (40,000 images), holdout set (10,000 images), and test set (20,000 images). Variants of the original MNIST digits dataset were constructed preserving the train/holdout/test split. For each variant, two datasets of equal size were constructed and denoted as primary and secondary (Supplementary Table 2).

For the ‘subset’ variant, the primary dataset was created by selecting half of the original images uniformly at random, and the secondary dataset comprised the complementary set of images. Thus, each subset contained half of the original data, and the two datasets together were equivalent to the original data.

For the ‘horizontal’ and ‘vertical’ variants, primary datasets were created by masking features corresponding to one side of the images. The secondary datasets were created by masking the opposite sides. In these variants, the primary and secondary datasets were jointly equivalent to the original data.

For variants denoted as ‘dropout’, ‘noise’, ‘erased’, and ‘spot’, the primary datasets consisted of all the images, but with some pixel intensities replaced by new values. ‘Dropout’ and ‘noise’ datasets selected half of the pixels at random and replaced their intensities by zero and nonzero values, respectively. ‘Erased’ and ‘spot’ corrupted circular regions with an area corresponding to half the number of active pixels. The secondary datasets were created using the same procedure, but using a different selection of pixels and circular regions. As a result of randomness, some pixel values remained the same in both the primary and secondary datasets, and some pixels were corrupted in both.

Datasets denoted as ‘missing’ and ‘lost’ were generated similarly to ‘dropout’ and ‘erased’, but values were set to not-available. Secondary datasets were generated by repeating the process. Thus, some pixels remained intact in both the primary and secondary datasets, and some pixels were corrupted in both.

One dataset variant denoted as ‘mislabel’ was generated by shuffling half of the image labels, effectively annotating images with erroneous digits. The primary and secondary datasets were generated with the same process with different selections of images to mislabel and different shuffling of the labels. Some labels remained accurate in both primary and secondary datasets, some were accurate in only one, and others were corrupted in both.

The ‘ramp’ variant was generated by selecting half of the images from the original datasets with a probability distribution that favored low digits. The secondary dataset was defined as the complementary set of images, and thus had an abundance of high-valued digits.

Separately to the generation of primary and secondary datasets for MNIST variants, two further dataset series were created as series. One series created a complete set of MNIST digit images with increasing proportions of missing pixels, from 0% to 80% in increments of 10%. Another series was created by adding random intensity values to increasing proportions of pixels, from 0% to 100% in increments of 10%.

### Machine learning with MNIST digits

All experiments with the MNIST digits dataset and its variants were performed using the xgboost library (Chen et al. 2019). All evaluations were performed using the hit rate as a metric, also called recall, which measures the proportion of images for which a model predicts the correct digit label.

An initial explorative analysis was performed on the original dataset and with the ‘subset’ variants. Multiclass classifiers were trained on the training tranches using arguments `objective=“multi:softmax”, num_class=10`, and probing the hyperparameter `nrounds` in the range from 2 to 50. All other parameters were kept at their defaults. Models were evaluated on training and holdout data (not used in training). As is common in ML training, there was a gap between performance on the two tranches, but a consistent increase in performance throughout this range indicated net benefits to longer training with higher values of `nrounds` (Supplementary Figure 1). An arbitrary value of `nrounds=30` was chosen for all subsequent experiments in order to limit processing time while achieving more than 96% recall.

Separate models were trained for the primary and secondary datasets of the MNIST variants. All models were trained only on the training tranches of their datasets. They were evaluated on the training and test tranches of their datasets, and also on the test tranche of the original images. For the ‘subset’ variant, evaluation on the original test images implies that the model trained on the primary dataset was evaluated on test images assigned to both the primary and secondary test tranche, and similarly for the model trained on the secondary tranche. For the other variants, the images in the variant-specific test tranche and the original test tranche differed in content. For the ‘missing’ variant, for example, the variant-specific test tranche consisted of images with missing values, whereas the test tranche of the original dataset consisted of the same images but with all the pixel intensities intact.

Ensembles were created by composing the primary and secondary models. Ensembles were evaluated on the test tranche of the original dataset. Uncalibrated ensembles averaged predictions produced by the two models using a simple mean (of softmax probabilities output by the xgboost models). Ensemble calibration was carried out using the holdout tranche of the original data (not used in training or for testing).

### Single-cell expression in testis

Datasets for two single-cell transcriptomics studies (Sohni et al. 2019; Guo et al. 2020) were obtained through the Single-cell Expression Atlas (Papatheodorou et al. 2020). Downloaded data included normalized gene-expression values in units of transcripts per million sequenced reads (TPM) and tables with cell-level annotations.

Marker genes for testis cell types were obtained from the first transcriptomic study (Sohni et al. 2019). Gene symbols were converted into Ensemble gene identifiers and trimmed to a set also included in the data obtained from the Single-cell Expression Atlas. Although the markers and data originated from the same study, such harmonization can lead to gene dropout due to mismatches in gene identifier-symbol nomenclature. In this case, 103 genes were linked to 16 different cell types.

Expression data for the first study were truncated to the 103 marker genes and log2-transformed with a unit pseudocount, i.e. values were replaced as *x* → log_2_(1 + *x*). A numerical value was computed for each cell-type signature as the mean of log2-transformed values for the marker genes for that signature.

Uniform manifold approximation and projection (Becht et al. 2019) implemented in R (Konopka 2020) was used with default settings to create visualizations in two dimensions. Cells in the embeddings were labeled with one label, defined as the cell type with the largest gene signature value defined above.

Expression data for the second study were truncated to the same 103 gene set and log2-transformed with a unit pseudocount. Cell-type labels were available for 7,416 out of a total of 31,565 cells, and subsequent analysis was restricted to the smaller set. This dataset, containing cells from four donors aged between 7 and 14 years old, was further split into parts for training, holdout, and testing (Supplementary Table 3). Labels were available for Sertoli cells, Leydig and peritubular myoid cells grouped together, and endothelial cells; other cell types were also labeled but represented a minority of cells, so they were grouped together as ‘other’ cells. These labels were treated as the ground truth.

A gene signature-based classifier was defined as a function computing the mean value for marker genes for the 16 cell types defined in the first study, and calling a cell type based on the dominant gene signature. These predictions were grouped into four classes to correspond to the available ground-truth annotations: Sertoli cells, Leydig and peritubular myoid cells together, endothelial cells, and other cells.

A dedicated multiclass classifier was trained using the training portion of the dataset and using the same 103 gene features as used to compute cell-type signatures. This classifier was trained using xgboost without tuning, setting the hyperparameter to an arbitrary `nrounds=10`. Other parameters were left at their defaults.

The ensemble was defined as a combination of the gene signature-based classifier and the gene expression multiclass xgboost model. All calibrations were performed using holdout data, either using cells from all donors, or using data specific to individual donors. All evaluations were performed using the test tranche.

## Availability

The R package ‘mlensemble’ is available at https://github.com/tkonopka/mlensemble. Scripts for analyses of imaging and genomic datasets are available at https://github.com/tkonopka/correcting-ml.

## Supplementary Figures

**Supplementary Figure 1.**
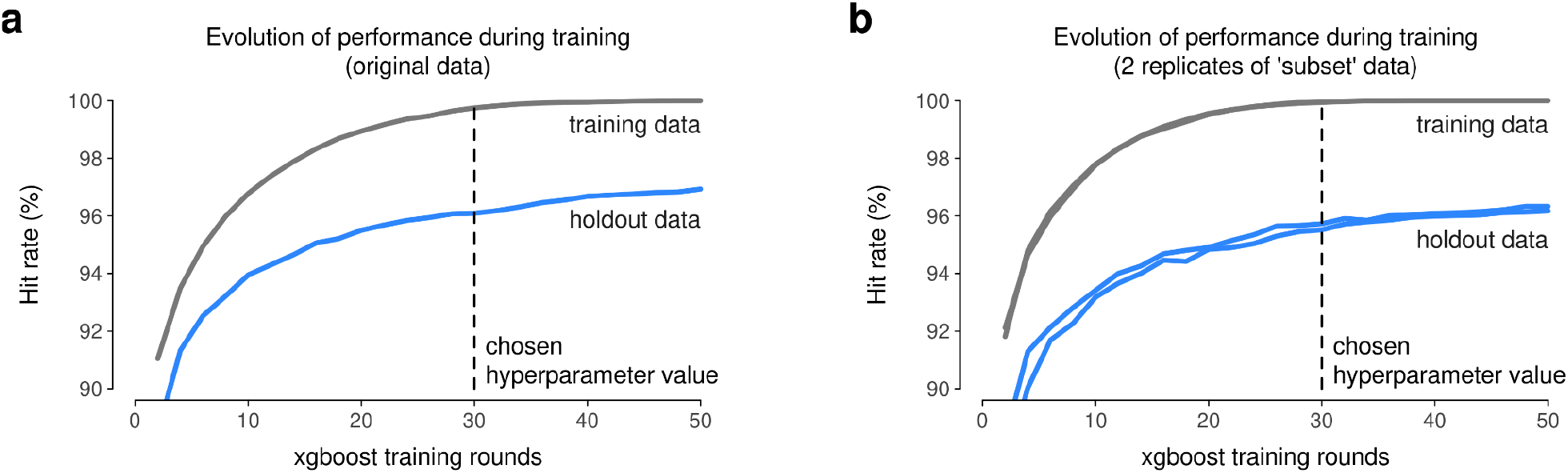
Training multiclass classifiers on the MNIST digits dataset and variants. **(a)** Performance of xgboost classification models trained and evaluated on the MNIST digits dataset. Performance is presented as a function of hyperparameter ’nrounds’, which determines the length of training. Performance is evaluated on the training data, which is available to the learning algorithm during training, and on holdout data, which is not available during learning. The vertical line indicates the hyperparameter value chosen for subsequent experiments because it lies along the upward-trending part of the performance curves and because it limits training time to an acceptable level for high-volume experiments. **(b)** Analogous to (a), but with calculations performed on two variants of the MNIST digits dataset, denoted as ‘subset’ variants, which each consist of half of the data instances as in the original dataset.

**Supplementary Figure 2.**
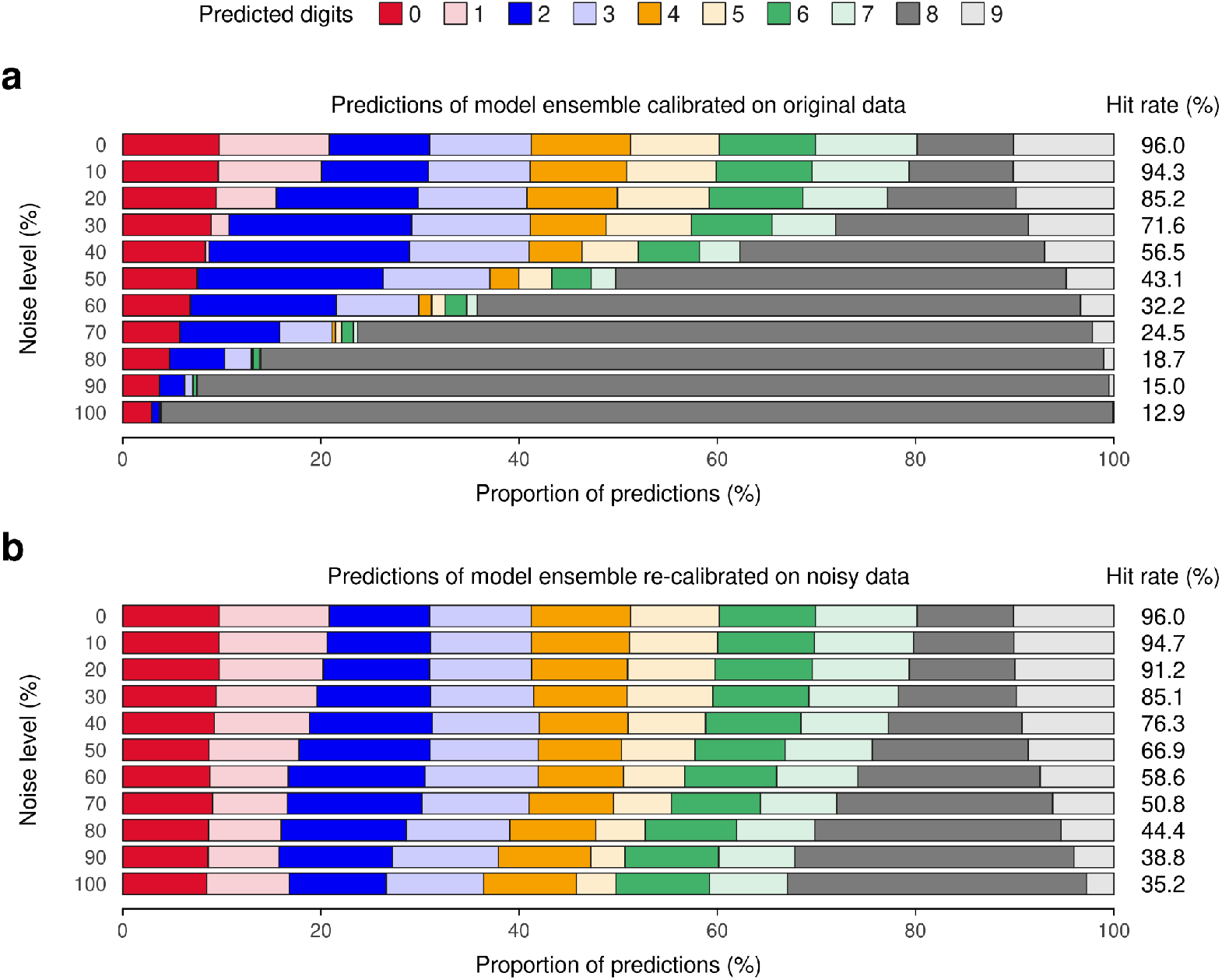
Predictions of model ensembles on MNIST digits with noise. **(a)** Breakdown of class predictions from a model ensemble applied to MNIST digits images with increasing levels of noise. The ensemble comprised two models, each trained on a distinct subset of MNIST images (without noise). The ensemble was calibrated on holdout images (without noise), and then applied on test images with increasing levels of noise. The hit rate on the right-hand side summarizes overall performance. **(b)** Analogous to (a), but with the model ensemble re-calibrated at each noise level on holdout images with noise.

## Supplementary Tables

**Supplementary Table 1.** Code examples using the `mlensemble` package. These examples are intended for illustration only and are not meant to be executable; working examples are available in the package documentation and the package vignette, which also introduce additional function parameters and package features.

| Analysis stage | Code examples |
| --- | --- |
| Model building | `# two models defined in base R # (assumes a data frame 'data_train') lm_1 <- lm(y ~ x1, data=data_train) lm_2 <- lm(y ~ x2, data=data_train) # two ml_model objects that wrap existing models model_1 <- ml_model(lm_1) model_2 <- ml_model(lm_2)` |
| Ensemble building | `ensemble <- model_1 + model_2` |
| Calibration | `# (assumes a data frame 'data_calib') ensemble_calibrated <- calibrate(ensemble, data=data_calib, label=data_calib$y)` |
| Prediction | `# (assumes a data frame 'data_test') predict(ensemble, data_test) predict(ensemble_calibrated, data_test)` |

**Supplementary Table 2.** MNIST dataset variants used for explorations of multiclass classification.

| **MNIST variant** | **Primary dataset** | **Secondary dataset** |
| --- | --- | --- |
| subset | ● half of the original instances selected at random;<br>● no adjustment to image pixels | ● complementary set of instances;<br>● no adjustment to image pixels |
| horizontal | ● all instances;<br>● features corresponding to right-hand side of images removed | ● all instances;<br>● features corresponding to left-hand side of images removed |
| vertical | ● all instances;<br>● features corresponding to bottom of images removed | ● all instances;<br>● features corresponding to top of images removed |
| dropout | ● all instances;<br>● half of pixels set to zero intensity | ● all instances;<br>● same procedure as for primary dataset, but using an independent sampling |
| noise | ● all instances;<br>● half of pixels set to a nonzero intensity | ● all instances;<br>● same procedure as for primary dataset, but using an independent sampling |
| erased | ● all instances;<br>● circular region set to zero intensity; circular region centered on a randomly selected pixel | ● all instances;<br>● same procedure as for primary dataset, but using an independent sampling |
| spot | ● all instances;<br>● circular region set to nonzero intensity; circular region centered on a randomly selected pixel | ● all instances;<br>● same procedure as for primary dataset, but using an independent sampling |
| missing | ● all instances;<br>● half of pixels set to missing (not available) values; missing pixels selected at random in each image | ● all instances;<br>● same procedure as for primary dataset, but using an independent sampling |
| lost | ● all instances;<br>● circular region set to missing (not available) values; circular region centered on a randomly selected pixel | ● all instances;<br>● same procedure as for primary dataset, but using an independent sampling |
| mislabel | ● all instances;<br>● half of labels shuffled | ● all instances;<br>● half of labels shuffled |
| ramp | ● half of instances; selected with a linear ramp probability distribution so that the 0 digit appears most often and the 9 digit least often<br>● no adjustment to pixel data | ● Complementary set of images compared to primary dataset<br>● no adjustment to pixel data |

**Supplementary Table 3.** Dataset sizes used in single-cell cell type classification experiments.

| Donor age | Training set (cells) | Holdout set (cells) | Test set (cells) |
| --- | --- | --- | --- |
| 7 years | 972 | 260 | 591 |
| 11 years | 460 | 118 | 259 |
| 13 years | 1260 | 364 | 719 |
| 14 years | 1296 | 386 | 728 |

